# Identification and characterization of a TRF2-like telomere-binding protein in Arabidopsis

**DOI:** 10.64898/2026.01.29.702503

**Authors:** Jana Fulneckova, Jana Faturova, Jaroslav Fulnecek, Jana Pecinková, David Potesil, Ganji Sri Ranjani, Zbynek Zdrahal, Karel Riha

## Abstract

Telomere protection and maintenance are mediated by proteins that bind telomeric DNA and recruit additional components of telomeric chromatin. While these factors are well characterized in yeast and mammals, their counterparts in plants remain poorly defined. Here, we used a proteomic approach in *Arabidopsis thaliana* to identify nuclear proteins that preferentially associate with telomeric DNA. We identified TRFL7, a previously uncharacterized member of the TRF-like (TRFL) protein family, as a prominent candidate. We show that TRFL7, together with its close homologues TRFL5 and TRFL11, associates with telomeric chromatin and forms distinct nuclear foci that preferentially localize near the nucleolus, resembling the nucleolus-associated telomere clustering characteristic of Arabidopsis. Genetic inactivation of TRFL7 in combination with either TRFL5 or TRFL11 results in telomere elongation, indicating a role for these proteins in telomere length homeostasis. Notably, TRFL7 contains an iDDR sequence motif that is also present in human TRF2, where it limits the activity of the Mre11-Rad50-Nbs1 complex. Together, our findings identify TRFL7 as a functional component of plant telomeric chromatin and suggest that it represents a plant orthologue of human TRF2.

## Introduction

Telomeres are specialized nucleoprotein structures that cap the ends of linear eukaryotic chromosomes, safeguarding genome integrity by preventing chromosome end–to–end fusions and inappropriate activation of DNA damage responses. They consist of tandem arrays of short, G-rich DNA repeats and an associated set of telomere-binding proteins that together form a protective cap and regulate telomere maintenance. Because conventional DNA polymerases cannot fully replicate chromosome termini, telomeres progressively shorten with each round of DNA replication. This shortening is counteracted by telomerase, which adds new telomeric repeats through a highly regulated process that maintains telomeres at an organism-specific length homeostasis.

Telomere-binding proteins play a central role in protecting chromosome ends and telomere maintenance. In vertebrates, a key component of telomeric chromatin is the shelterin complex, which consists of six proteins, the double-stranded telomeric DNA–binding proteins TRF1 and TRF2, POT1 that binds to the single-stranded 3’telomeric overhang, and the adaptor proteins TIN2, TPP1, and RAP1, which link these components into an integrated complex (1,2). POT1 and the TRF proteins play complementary roles in telomere protection and regulation. POT1 prevents activation of the ATR-mediated DNA damage response and regulates telomerase access through its interaction with TPP1 (3,4). In contrast, TRF1 and TRF2 bind double-stranded telomeric DNA and are essential for maintaining telomere structure. TRF2 promotes formation and stabilization of the protective T-loop and suppresses ATM signaling and end-to-end chromosome fusions (5,6), while TRF1 facilitates efficient telomere replication by preventing replication stress at telomeric repeats (7).

A shelterin-like complex is also present in fission yeast, exhibiting similar functionality and architecture to the human shelterin complex. It consists of the duplex telomere-binding protein Taz1, a functional homologue of TRF1 and TRF2, which is linked to the single-stranded telomere-binding protein Pot1 via the adaptor proteins Rap1, Poz1, and Tpz1 (8-10). The similarity of shelterin complexes suggests that the core principles of telomere protection and length regulation are evolutionarily conserved. However, this view is challenged by observations in budding yeast and Drosophila, which have more derived telomere architectures and lack a shelterin complex analogous to those found in mammals and fission yeast. This highlights the evolutionary flexibility of telomere maintenance strategies (11-13).

Plants offer an interesting perspective on the evolution and core mechanisms of telomere function. Although they diverged early from the animal–fungal lineage, they nevertheless share many fundamental features of telomere biology with vertebrates, including developmentally regulated telomerase, POT1 proteins, the CST complex, and the ability to form T⍰loops, structures that likely do not occur in budding or fission yeast (11,14,15). At the same time, plant telomeres display distinctive characteristics. Notably, a subset of plant chromosome ends lacks a 3⍰G⍰overhang, a feature that may reflect differences in how leading- and lagging⍰strand replication processes shape telomere end structure (16). These combined ancestral and lineage⍰specific features make plants a valuable system for understanding both the conserved and divergent mechanisms that govern telomere protection and maintenance across eukaryotes.

Research conducted over the past 30 years has yielded important insights into the organization and function of plant telomeres, including the composition and evolution of telomerase and telomeric DNA (17,18), the discovery of the CST complex (19,20), the characterization of plant POT1 orthologues (21- 23), and the elucidation of the role of Ku in telomere protection (16,24). Nonetheless, the full complement of proteins that bind duplex telomeric DNA in plants has not yet been identified. Initial attempts to isolate proteins capable of binding single- and double-stranded plant telomeric sequences led to the identification of several candidate proteins in *Arabidopsis*, rice, and tobacco (25-28). Some of these proteins contain a single SANT/Myb domain homologous to the Myb domain in human TRF1 and TRF2 that harbors an evolutionarily conserved telobox motif responsible for sequence-specific binding to telomeric DNA (29).

Systematic screens revealed three classes of telobox-containing proteins in plant genomes. Proteins of the TRB/SMH family exhibit tripartite structure with N-terminal SANT/Myb domain, globular histone H1- like domain, and C-terminal coiled-coil region and specifically bind to telomeric DNA *in vitro* (30,31). Extensive functional characterizations in Arabidopsis revealed that TRBs act as regulators of gene expression through recruiting chromatin modifying complexes to promoters that contain a short telomeric motif (32-35). TRBs can also associate with plant telomeres (36), and simultaneous inactivation of multiple TRB paralogues in Arabidopsis and moss *Physcomitrium* patens leads to shorter telomeres (32,37). Nevertheless, growth phenotypes reported in the trb1/2/3 mutants do not appear to be linked to telomeres, but rather to the deregulation of genes repressed by the PRC2 complex (32).

Another class of plant telobox-containing proteins carries a C-terminal SANT/Myb domain, as in human TRF1 and TRF2, and these proteins were termed TRFLs (TRF-like) (38). TRFL proteins are further subdivided into two families. Members of TRFL Family 1 form homodimers and bind telomeric DNA *in*. They are characterized by an extension of the SANT/Myb domain that forms an additional α-helix, contributing to the structural integrity of the canonical three-helix SANT/Myb fold (39,40). While mutation of the rice RTBP1 leads to telomere elongation and genome instability (41), simultaneous mutation of all TRFL Family 1 members in Arabidopsis does not result in a discernible impact on telomere length homeostasis (42). This suggests that TRFL Family 1 proteins do not play essential roles at Arabidopsis telomeres. TRFL Family 2 comprises an evolutionarily heterogeneous group of proteins with little sequence homology beyond the SANT/Myb domain. These proteins were reported to fail to interact with telomeric DNA *in vitro* (38), and this family has not yet been extensively characterized *in vivo*.

The absence or relatively mild telomeric phenotypes in plants deficient in the telobox proteins characterized so far indicates that our understanding of plant telomere-binding factors is incomplete. To address this gap, we performed a proteomic analysis of proteins that specifically bind to telomeric DNA in Arabidopsis. This approach led to the identification of a subgroup of telobox proteins belonging to TRFL Family 2. Functional characterization of these proteins suggests that they are components of Arabidopsis telomeres involved in regulating telomere length homeostasis, and are evolutionarily related to the human TRF2 protein.

## Material and methods

### Plant material and growth conditions

*Arabidodpsis thaliana* (ecotype Col-0) plants were grown in phytotrons under LED illumination (white 77%/red 20%/infrared 3%; 150 μmol m^−2^ s^−1^) with 16/8-h light/dark regime. The T-DNA insertion lines used in the study were obtained from the Nottingham Arabidopsis Stock Centre: *trfl5* (SAIL_504G05), *trfl7* (SAIL_1155F11), *trfl11*-1 (SALK_053119c) and *trfl11*-2 (SALK_018118c). PCR primers used for genotyping the mutant lines are listed in Table S1.

### Preparation of nuclear protein extracts

Nuclei were isolated from protoplasts prepared from an A. thaliana root suspension culture. Cells from 150 ml of culture were harvested by centrifugation (310 × g, 5 min). For enzymatic digestion, 5 ml of pellet was mixed with 25 ml of enzyme solution containing 0.6% cellulase Onozuka R-10 (Ducheva) and 0.2% Macerozyme R-10 (Ducheva) in GM solution (0.34 M glucose, 0.34 M mannitol, 4.4 g L^−1^ Murashige & Skoog medium with vitamins, pH 5.5), and the volume was adjusted to 50 ml. The suspension was incubated in the dark with gentle agitation for 5 h. Protoplasts were collected by centrifugation (200 × g, 5 min, 4 °C), washed in GM solution, and resuspended in S-solution (0.28 M sucrose, 4.4 g L^−1^ Murashige & Skoog medium with vitamins, pH 5.5). Floating protoplasts were recovered by centrifugation (130 × g, 10 min, 4 °C), adjusted to 50 ml with GM solution, and pelleted (100 × g, 5 min, 4 °C).

Nuclei were released by adding 5 volumes of hypotonic lysis buffer (10 mM HEPES pH 7.9, 1.5 mM MgCl_2_, 10 mM KCl, 0.5 mM DTT, 0.05% NP-40) to 1 volume of protoplast suspension and incubating on ice for 15 min. The lysate was filtered through 70 µm and 40 µm meshes and Miracloth, mixed with 100 ml Nuclei Isolation Buffer (NIB: 10 mM MES pH 6.0, 10 mM NaCl, 5 mM EDTA, 250 mM sucrose, 20 mM β-mercaptoethanol, 150 µM spermidine, 100 µM PMSF) containing 0.1% Triton X-100, and centrifuged (770 × g, 15 min, 4 °C). The pellet was resuspended in the same buffer, incubated on ice for 15 min, and centrifuged again. Nuclei were purified by resuspension in 5 ml buffer E (7.5 g Percoll, 1 g 5× NIB) and centrifugation (5900 × g, 10 min, 4 °C). Floating nuclei were washed twice in NIB (770 × g, 15 min, 4 °C) to remove Percoll.

For protein extraction, nuclei were mixed with 2 volumes of Nuclear Lysis Buffer (1.2 M KCl, 20 mM HEPES pH 7.9, 25% glycerol, 2 mM MgCl_2_, 0.2 mM EDTA, 0.1% NP-40, 2 mM DTT, protease inhibitors) and incubated for 30 min at 4 °C. After centrifugation (30 000 × g, 20 min, 4 °C), supernatants were aliquoted, stored at −80 °C, and protein concentrations determined by Bradford assay (Bio-Rad).

### Affinity purification of telomeric-DNA binding proteins

The 50 bp probes were prepared by annealing end-biotinylated oligonucleotides (Table S1). The 200 bp baits were generated by non-template PCR containing 200 µM each dNTP, 2 µM biotinylated G-strand oligonucleotide, 2 µM complementary C-strand oligonucleotide (Table S1), and 2 U KAPA Taq polymerase (Riche) in 1× Buffer A. Following initial denaturation (95 °C, 3 min), amplification consisted of 3 cycles of 95 °C for 1 min, 55 °C for 30 s, and 72 °C for 1 min, followed by 40 cycles of 95 °C for 1 min, 60 °C for 30 s, and 72 °C for 20 s, with a final extension at 72 °C for 5 min. PCR products were verified by agarose gel electrophoresis and purified using the NucleoSpin™ Gel and PCR Clean-up Kit (Macherey-Nagel).

For pull-down assays, 250 µl of nuclear extract (170 µg total protein) was incubated with 500 µg Dynabeads™ M-280 Streptavidin (Invitrogen) bearing 5 µg of telomeric or control DNA. As a negative control, nuclear extract was incubated with beads lacking immobilized DNA. The mixtures were placed in dialysis tubing and incubated overnight at 4 °C in DB buffer (20 mM HEPES, pH 7.9; 20% glycerol; 20 mM KCl; 1.5 mM MgCl_2_; 0.2 mM EDTA; 0.2 mM PMSF; 2 mM DTT) to allow protein binding. Beads were washed four times for 5 min with rotation in 400 µl PBB buffer (50 mM Tris–HCl, pH 8.0; 150 mM NaCl; 5 mM MgCl_2_; 0.5% NP-40; 1 mM DTT; Complete Protease Inhibitor without EDTA [Roche]), followed by two washes with PBS, and finally resuspended in 20 µl PBS.

### Mass spectrometry

Proteins were extracted from beads with 2% SDS in a thermomixer (50°C, 750 rpm, 20 min), then centrifuged (2 min, 1,000 × g) and the supernatant transferred to a clean tube. Dithiothreitol (0.5 M stock) was added to a final concentration of 0.1 M, and proteins were reduced at 95°C for 15 min. Samples were processed by filter-aided sample preparation (FASP) (43) using 0.75 µg trypsin (18 h, 37°C; sequencing grade, Promega). Resulting peptides were cleaned by three rounds of liquid–liquid extraction with water-saturated ethyl acetate (44), dried in a SpeedVac, and reconstituted in 2.5% formic acid in 50% acetonitrile and 100% acetonitrile containing 0.001% polyethylene glycol, then concentrated in a SpeedVac.

LC–MS/MS analyses were performed on an Ultimate 3000 RSLCnano system coupled to an Orbitrap Fusion Lumos mass spectrometer (Thermo Fisher Scientific). Tryptic digests were online concentrated and desalted on a trapping column (300 µm × 5 mm, Acclaim PepMap100 C18, 5 µm; Thermo Fisher Scientific) and eluted at 300 nL/min onto an analytical Aurora C18 column (75 µm × 250 mm, 1.6 µm; IonOpticks) using an 86 min linear gradient (3–37% B; A: 0.1% FA in water, B: 0.1% FA in 80% ACN). Columns were equilibrated before injection, the analytical column was maintained at 50°C, and spray voltage and sheath gas were set to 1.5 kV and 1, respectively.

Data were acquired in DIA mode with MS1 scans from m/z 350–1650 at 60,000 resolution (AGC 300%, 55 ms) and HCD MS2 scans at 27% NCE from m/z 200–1800 at 30,000 resolution (AGC 1000%, 55 ms). Staggered isolation windows (20 Th) covered m/z 400–1000. Raw files were converted to mzML using MSConvertGUI (v3.0.21193) with vendor peak picking (MS1) and demultiplexing (10 ppm mass error). mzML files were processed in DIA-NN (v1.8)(45) in library-free mode against the cRAP database (111 entries) and the *Arabidopsis thaliana* UniProtKB proteome (27,474 sequences, 2022-06-16). Carbamidomethylation was set as a fixed modification, trypsin/P with one missed cleavage and peptide lengths of 7–30 aa were allowed, and FDR was controlled at 1%. MS1 and MS2 mass tolerances were set to 6 and 16 ppm, respectively, with match-between-runs enabled. Protein MaxLFQ intensities were processed using OmicsWorkflows (v4.1.3a), including removal of low-quality precursors and contaminants, filtering for proteotypic peptides, log2 transformation, replicate-based filtering, and differential expression analysis using LIMMA. Quantitative candidates were selected with |log2FC| > 1 and adjusted P < 0.05, and qualitative changes were defined as proteins detected in >50% of positive samples and absent in all nonTelo controls.

### Protein-DNA binding assay

cDNA fragments encoding full-length TRFL7 and a TRFL7 deletion variant lacking the MYB domain were amplified from Arabidopsis cDNA using Phusion™ Hot Start II DNA Polymerase (Thermo Fisher Scientific) and the primer pairs TRFL7-BamHI Fw/TRFL7-XhoI R and TRFL7-BamHI Fw/TRFL7-XhoI-Myb R, respectively (Table S3). PCR products were cloned into the pJET1.2/blunt vector using the CloneJET PCR Cloning Kit (Thermo Fisher Scientific) and subsequently subcloned into the SspI site of the 1M vector (pET His_6_-MBP-TEV LIC vector) by ligation-independent cloning (LIC) using primer pairs TRFL7-LIC-F/TRFL7-LIC-R (full-length) and TRFL7-LIC-F/TRFL7-MYB-LIC-R (ΔMYB variant) (Table S3).

Proteins were expressed in Escherichia coli Rosetta (DE3) by induction with 0.2 mM IPTG at 30 °C for 3 h. Cells were harvested (4,400 × g, 15 min, 4 °C) and resuspended in lysis buffer (0.1 M Tris–HCl, pH 7.5; 0.5 M NaCl; 0.05% Tween 20; 1% Triton X-100; 1 mg mL^−1^ lysozyme; 1 mM DTT; 1× protease inhibitor cocktail [EDTA-free; Roche]). The suspension was sonicated and centrifuged (14,000 × g, 20 min, 4 °C), and the supernatant was filtered through a 0.2 µm membrane. Imidazole was added to 50 mM, and His-tagged proteins were purified using His Mag Sepharose™ Ni magnetic beads (Cytiva) according to the manufacturer’s instructions. Proteins were eluted in buffer containing 0.1 M Tris–HCl (pH 7.5), 0.5 M NaCl, and 0.5 M imidazole.

For pull-down assays, 5 µg biotinylated DNA probes were immobilized on 500 µg Dynabeads™ M-280 Streptavidin (Invitrogen) and incubated for 1 h at 4 °C with either 11 µg purified recombinant TRFL7 or TRFL7 ΔMYB protein, or 10 µl crude E. coli lysate, in binding buffer (25 mM HEPES, pH 7.5; 150 mM KCl; 5 mM MgCl_2_; 10% glycerol; 1 mM DTT; 1× EDTA-free protease inhibitor cocktail [Roche]). As a negative control, proteins were incubated with beads lacking bait DNA. Beads were washed five times with PBB buffer (50 mM Tris–HCl, pH 7.5; 150 mM NaCl; 5 mM MgCl_2_; 0.5% NP-40; 1 mM DTT; EDTA-free protease inhibitors), and bound proteins were eluted by heating in SDS loading buffer at 85 °C for 10 min. Proteins were resolved by 12% SDS–PAGE, transferred to a PVDF membrane (Thermo Scientific), and visualized by Ponceau S staining. His-tagged proteins were detected using a mouse monoclonal anti-6×His antibody (Abcam, ab18184) followed by IRDye® 800CW goat anti-mouse IgG (LI-COR Biosciences).

### Phylogenetic analysis

FASTA protein sequences were retrieved from the PANTHER database and the NCBI protein database (Supplementary Dataset 1). Sequences were aligned using MUSCLE as implemented in MEGA 11 with default parameters (46). Phylogenetic analysis was performed in MEGA 11 using the maximum likelihood method with 1,000 bootstrap replicates (47). Human TRF1 and TRF2 protein sequences were used as outgroups to root the phylogenetic tree

### Protein localization

Total RNA was extracted from 7-day-old *A. thaliana* Col-0 seedlings using TRI Reagent (Molecular Research Center) according to the manufacturer’s instructions. Genomic DNA was removed with the TURBO DNA-free™ Kit (Invitrogen). First-strand cDNA was synthesized from 1 µg total RNA using RevertAid Reverse Transcriptase (Thermo Scientific).The coding sequences of TRFL3 (At1g17460), TRFL5 (At1g15720), TRFL6 (At1g72650), TRFL7 (At1g06910), TRFL8 (At2g37025), TRFL9 (At3g12560), TRFL10 (At5g03780), and TRFL11 (At5g58340) were amplified from cDNA using Q5 Hot Start High-Fidelity DNA Polymerase (New England Biolabs). PCR products were fused to Gateway® attB sites and cloned into the pDONR/Zeo entry vector using BP Clonase™ (Invitrogen), then sequence-verified. Primer sequences are listed in Table S1. Coding sequences were transferred into destination vectors pGWB641 (C-terminal YFP), pGWB642 (N-terminal YFP), pGWB460 (C-terminal RFP), or pGWB461 (N-terminal RFP) using LR Clonase™ II (Invitrogen) (48). Constructs were either transfected into leaf protoplasts as described previously (49) or introduced into Agrobacterium tumefaciens strain GV3101 for plant transformation by the floral dip method. Transformed material was imaged using a Zeiss LSM780 confocal microscope.

### Chromatin immunoprecipitation

Chromatin was isolated from 10-day-old seedlings and sheared as described previously (24). Prior to immunoprecipitation, the sheared chromatin was centrifuged twice (25,000 × g, 10 min, 4 °C) to remove debris. The clarified chromatin was incubated with GFP-Trap® Magnetic Agarose (Chromotek), while control samples were processed in parallel using magnetic agarose lacking the anti-GFP nanobody. Beads were equilibrated in IP buffer (50 mM HEPES, pH 7.5; 150 mM NaCl; 5 mM MgCl_2_; 10 µM ZnSO_4_; 0.1% Triton X-100) supplemented with 1% BSA before immunoprecipitation. Chromatin binding was performed at 4 °C for 30 min on a rotating wheel. Beads were washed five times with 1 ml IP buffer at 4 °C for 10 min per wash, transferring beads to new low-DNA-binding tubes after the second and fourth washes. After these washes, beads were resuspended in 150 µl TE–NaCl buffer (TE buffer supplemented with 0.5 M NaCl).

For decrosslinking, 150 µl TE–NaCl buffer was added to 130 µl of input sample, and all samples designated for decrosslinking were treated with 1 µl RNase A (10 mg ml^−1^) and incubated at 37 °C for 1 h. Proteinase K (2 µl, 20 mg ml^−1^) was then added, followed by incubation at 37 °C for 2 h and overnight incubation at 65 °C. The next day, samples were adjusted to 450 µl with TE buffer and extracted once with 500 µl phenol–chloroform–isoamyl alcohol (25:24:1) and once with 500 µl chloroform–isoamyl alcohol (24:1). DNA was precipitated by adding 1 µl glycogen (20 mg ml^−1^) and 3 volumes of ethanol, followed by centrifugation (25,000 × g, 30 min, 4 °C), washing with 750 µl 70% ethanol, air-drying, and resuspension in 50 µl 5 mM Tris–Cl (pH 8.0).

One microliter of DNA was used in a 20 µl quantitative PCR (qPCR) reaction containing 10 µl of 2× KAPA SYBR® Fast Universal Master Mix and 0.25 µM each of TelA and TelB primers (50) for telomeric DNA amplification, or CEN-f and CEN-r primers for centromeric DNA amplification. PCR conditions were as follows: 95 °C for 3 min, followed by 45 cycles of 95 °C for 10 s, 60 °C for 20 s, and 72 °C for 1 s, with fluorescence acquisition at the end of each cycle. Melting curve analysis was performed at the end of amplification (95 °C for 5 s, 65 °C for 1 min, and 97 °C for 1 s, with five acquisitions per °C). Absolute quantification was performed using a calibration curve generated from a series of 10-fold dilutions of input DNA to determine amplification efficiency. Target quantities were normalized either to input DNA or expressed as a ratio relative to the negative control, consisting of samples immunoprecipitated with control magnetic beads lacking the GFP nanobody.

### Yeast two-hybrid assay

Yeast two-hybrid assays were performed using the Matchmaker™ GAL4-based two-hybrid system (Clontech). The coding sequences of TRFL5, TRFL7, and TRFL11 were subcloned from their entry clones into the destination vectors pGADT7-DEST and pGBKT7-DEST using LR Clonase™ (Invitrogen). Each bait– prey combination was co-transformed into Saccharomyces cerevisiae strain PJ69-4a and grown at 30 °C on synthetic dropout (SD) agar lacking leucine and tryptophan (–Leu, –Trp; –LW) to select for co-transformants. Selected colonies were inoculated into YPD medium and grown overnight. Successful co-transformation was confirmed by growth on –LW medium, and protein–protein interactions were assessed on SD medium lacking leucine, tryptophan, and histidine (–Leu, –Trp, –His; –LWH). Interaction strength was evaluated by supplementing –LWH medium with increasing concentrations of 3- aminotriazole (3-AT). All experiments were performed in two independent biological replicates, each with three technical replicates.

### Terminal restriction fragment analysis

Genomic DNA was isolated from young rosette leaves using a standard phenol–chloroform extraction protocol. Terminal restriction fragment (TRF) analysis was performed using TruI restriction endonuclease (Thermo Scientific) as previously described (51).

## Results

### Identification of telomeric-DNA biding proteins from Arabidopsis nuclei

To identify proteins that bind telomeric double stranded DNA, we performed quantitative label-free mass spectrometry of nuclear proteins specifically enriched in pull-downs with telomeric DNA (52,53). Two types of telomeric probes, along with scrambled (TGTAGAG)_n_ controls, were used in parallel for the pull-down experiments (Figure 1A): 50 bp probes composed of two annealed oligonucleotides containing five TTTAGGG repeats, and 200 bp probes generated by non-template PCR using telomeric oligonucleotides. The pull-down experiments with 50 pb probes yielded 247 proteins that were significantly enriched in purifications with the TTTAGGG probe compared with the scrambled control (>2- fold enrichment; adjusted P < 0.05; Figure 1B). Gene ontology enrichment analysis for molecular function (54) revealed RNA and single-strand DNA bindings among the most enriched categories (Figure S1). The observed prevalence of single-stranded nucleic acids binding activities likely reflects affinity to non-annealed oligonucleotides that may have been attached to the beads.

**Figure 1.**
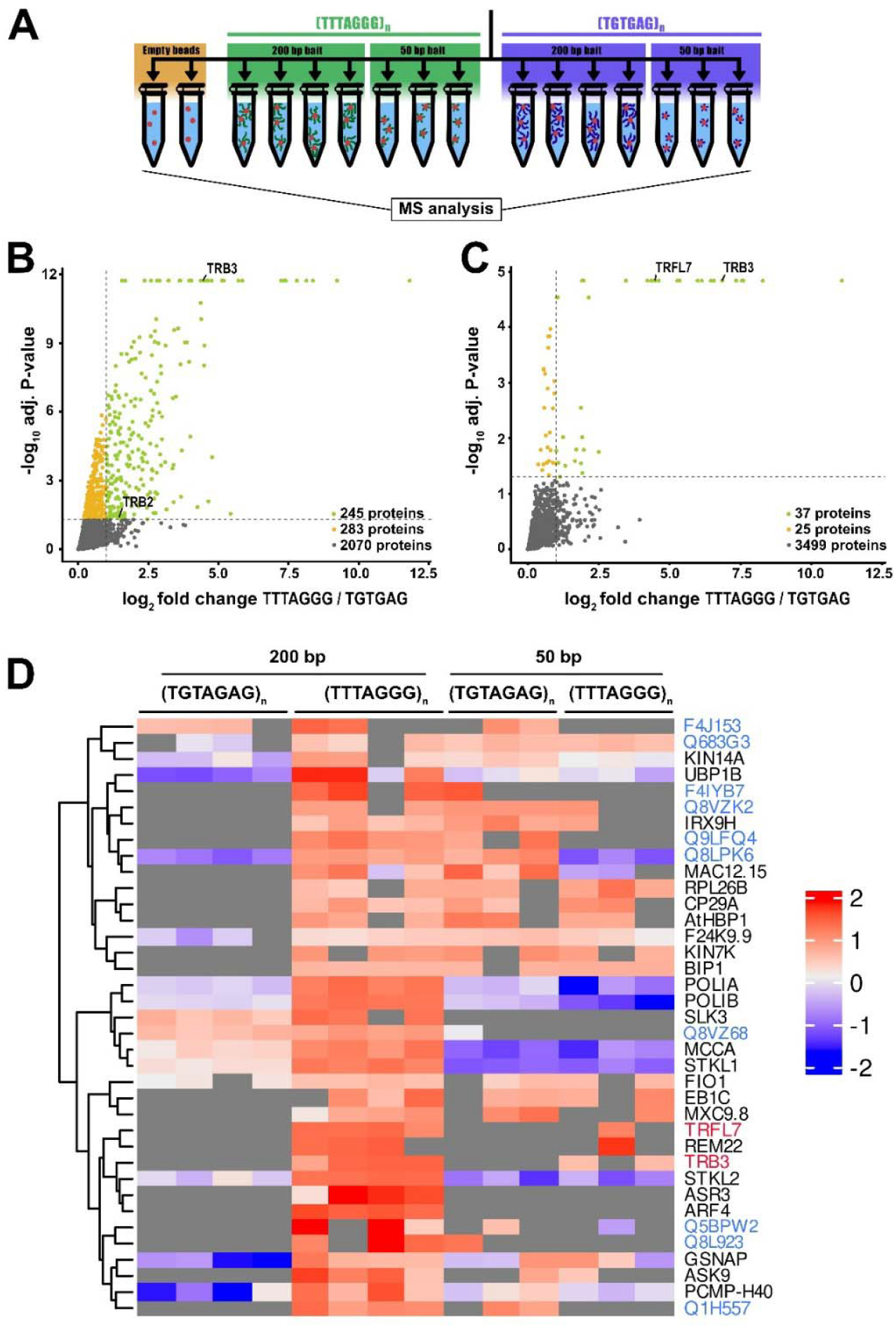
Identification of telomeric DNA-binding proteins. **(A)** Scheme of the telomeric DNA affinity purification experiment. (**B, C**) Volcano plots showing significantly enriched proteins in affinity purifications with 50 bp (B) and 200 bp (C) probes. Telobox-containing proteins are indicated. (**D**) Heat- map representation of z-score enrichment for proteins specifically enriched in the pull-downs with the telomeric 200 bp probe. Blue labels indicate UniProt protein identifiers, telobox-containing proteins are labeled in red.

In contrast, only 37 out of 3499 detected proteins showed significant enrichment in pull-downs with the 200 bp probe (Figure 1C,D). Among these were several transcription factors, including ARS3, ARF4, SLK3, REM22 and STKL1/2, indicating that the 200 bp probe conferred greater specificity toward double-strand DNA binding proteins. Importantly, pull-downs with both probes identified TRB3, a protein that has previously been shown to bind telomeric DNA and associate with telomeres (30,31), and that has also been detected in similar pull-down experiments (53). In addition, another telobox protein, TRFL7, was specifically enriched in all experiments with the 200 bp probe and in one pull-down experiment with the 50 bp probe; no enrichment was detected in experiments with scrambled probes (Figure 1C,D).

This observation was unexpected as TRFL7 belongs to the TRFL Family 2 proteins, and a previous study failed to detect an interaction between TRFL7 and telomeric DNA *in vitro* (38). To validate that the presence of TRFL7 in affinity purifications from Arabidopsis nuclear extracts was due to direct binding to telomeric DNA rather than indirect association through another protein, we repeated the pull-down experiments using proteins ectopically expressed in *Escherichia coli* (Figure 2). Because TRFL7 has a tendency to aggregate, it was expressed as a fusion with maltose-binding protein (MBP) to increase its solubility. His-MBP-TRFL7 specifically co-purified with the telomeric probe from both whole- cell lysates and affinity-purified protein fractions (Figure 2A). Importantly, this co-purification was substantially reduced in the presence of the scrambled probe or when using constructs lacking the C-terminal SANT/Myb domain (Figure 2A,B). This data indicated that TRFL7 specifically binds to telomeric DNA and that the interaction involves the SANT/Myb domain.

**Figure 2.**
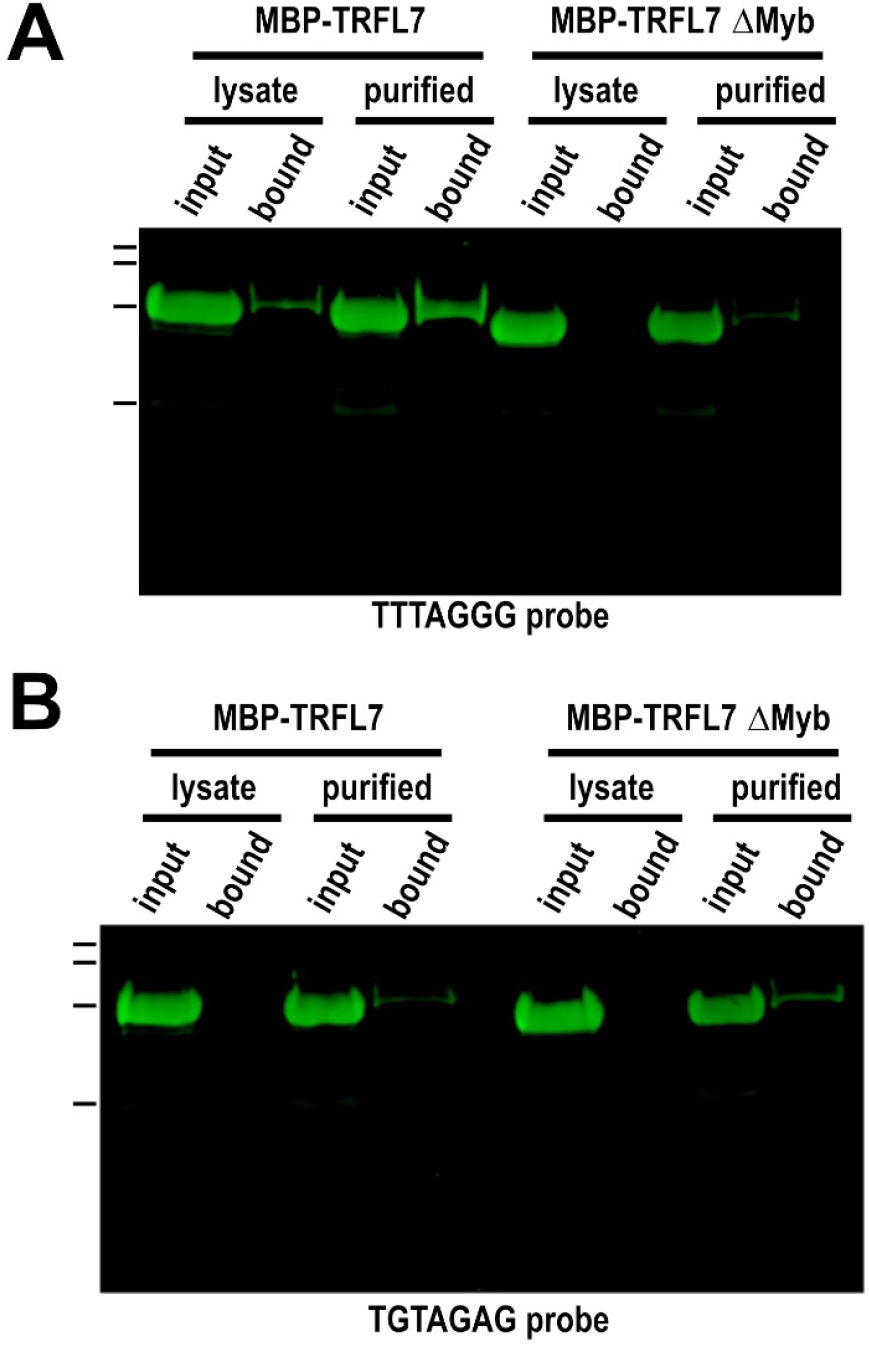
Co-purification of TRFL7 with telomeric DNA. Cell lysates from bacteria expressing His–MBP– TRFL7 or purified protein were incubated with 200-bp telomeric (**A**) or scrambled (**B**) DNA probes, followed by coprecipitation and visualization of captured proteins by Western blotting using an anti-His antibody.

### TRFL7 protein is phylogenetically related to human TRF2

Previous studies described seven TRFL Family 2 proteins in the *A. thaliana* genome: TRFL3, TRFL5, TRFL6, TRFL7, TRFL8, TRFL10 and TRFL11 (38,55). TRFL family 2 appears to be evolutionary more heterogeneous than the TRFL Family 1 (38). To gain deeper insight into the evolutionary relationships among TRFL proteins, we performed a phylogenetic reconstruction using genomes representing diverse phylogenetic groups of green plants (Figure 3A). *Klebsormidium nitens*, a green alga that diverged from land plants approximately 980-680 million years ago (56), already possess representatives of both TRFL families. This observation indicates that TRFL Family 1, characterized by the Myb extension domain, emerged at the base of green plant evolution.

**Figure 3.**
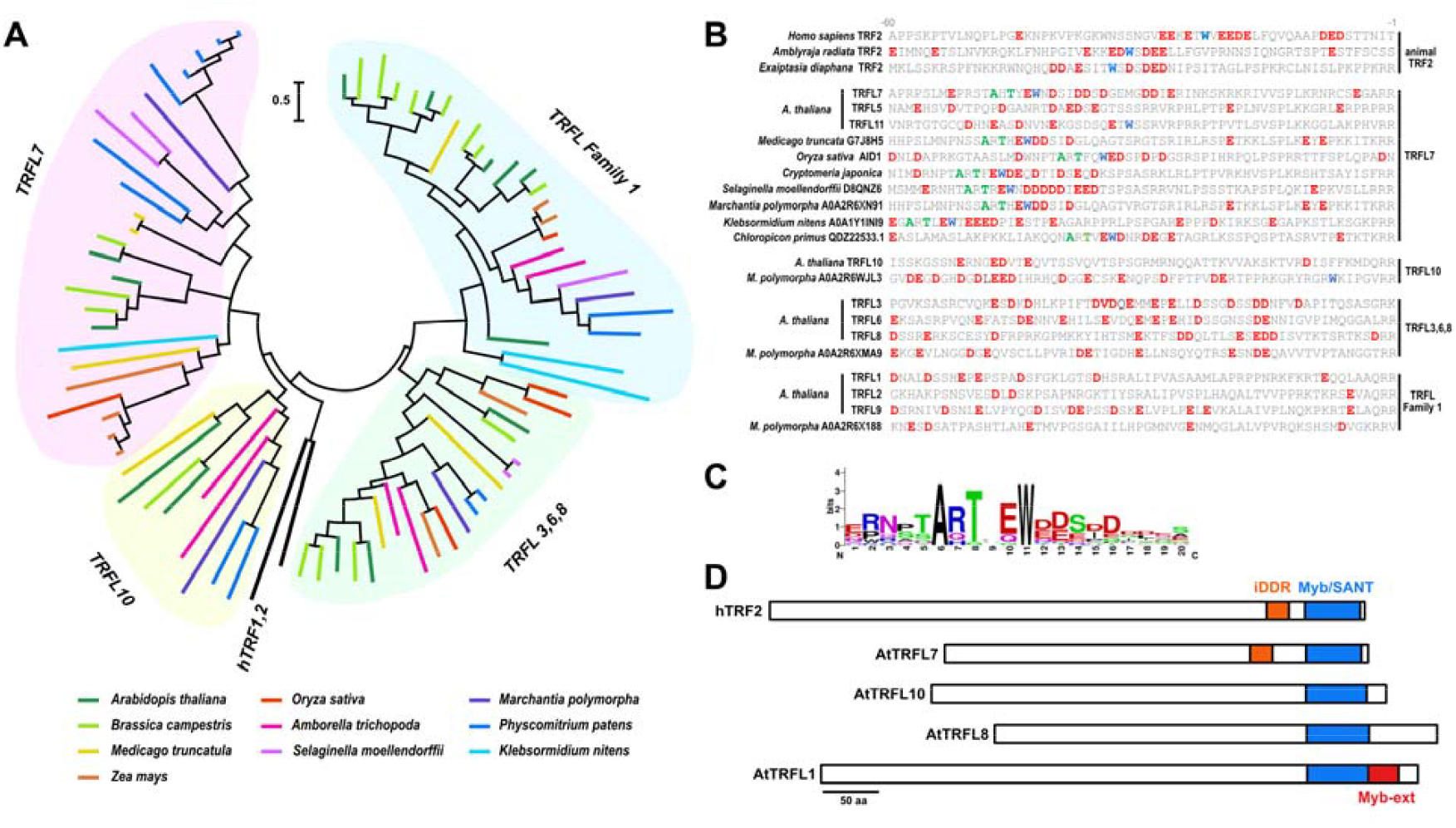
Phylogenetic analysis of plant TRFL proteins. **(A)** Phylogenetic tree of TRFL homologues from the indicated plant species. The scale bar indicates evolutionary distance. More detailed version of the tree with the bootsrap values is shown in Figure S2 (**B**) Comparison of the 60 amino acids immediately preceding the SANT/Myb domain in the indicated TRF2 and TRFL proteins. Acidic residues (red), the conserved tryptophan (blue) in the iDDR motif, and plant-specific alanine and threonine residues (green) are highlighted. (**C**) Sequence logo showing conserved residues within the iDDR motif of plant TRFL7 orthologues. (**D**) Domain topology of human TRF2 and Arabidopsis representatives of distinct TRFL subgroups, as classified in the phylogenetic tree.

Proteins of the TRFL Family 2 segregate into three distinct groups, which we designate according to their *A. thaliana* representatives as the TRFL7, TRFL10 and TRFL3,6,8 subgroups (Figure 3A). This diversification occurred at the base of land plants, as all three subgroups are present in the liverwort *Marchantia polymorpha* and the moss *P. patens*. The sole TRFL Family 2 orthologue of *K. nitens* belongs to the TRFL7 subgroup, suggesting that TRFL7 proteins have retained the ancestral structural features of this family. Notably, human hTRF1 and hTRF2, whose SANT/Myb domains are most closely related to TRFL1 Family 2 proteins (38), also tend to cluster with the TRFL7 and TRFL10 subgroups (Figure 3A, Figure S2).

Phylogenetic analysis in metazoans indicates that TRF2-like factors represent the original components of the shelterin complex and that TRF1 emerged from TRF2 via an ancient duplication event in the vertebrate lineage (57). *A defining* feature of metazoan TRF2-like proteins, in addition to the SANT/Myb domain, is the presence of the so called iDDR motif. This motif consists of a stretch of acidic amino acid residues flanking a tryptophan residue immediately upstream of the SANT/Myb domain and is required for attenuation of the DNA damage response mediated by ATM signaling (57,58).

Strikingly, inspection of the corresponding region in Arabidopsis TRFL7 revealed a similar stretch of aspartate and glutamate residues surrounding a conserved tryptophane (Figure 3B). Extension of this analysis to TRFL7 orthologues from other green plants identified a highly conserved sequence motif comprising invariant alanine and threonine residues followed by a tryptophan flanked by acidic amino acids (Figure 3C). This motif is present in TRFL7 homologues from all examined plant species, including the green algae *K. nitens* and *Chloropicon primus*, representatives of streptophyte and chlorophyte lineages, respectively, which diverged more than one billion years ago at the base of plant evolution. The deep evolutionary origin of the iDDR motif, together with the shared protein topology characterized by a telobox-containing C-terminal SANT/Myb domain immediately preceded by the iDDR motif (Figure 3D), strongly argues that plant TRFL7 and metazoan TRF2-like proteins originated from a common ancestral protein and have been constrained to retain these features.

The *Arabidopsis* TRFL7 subgroup consists of three proteins, TRFL7, TRFL5, and TRFL11, which originated from relatively recent duplication events within the Brassicaceae (Figure 3A; Figure S2). Despite their recent evolutionary origin, the iDDR motif has partially degenerated in TRFL5 and TRFL11 and is retained only in TRFL7. This observation suggests that TRFL5 and TRFL11 may have partially diverged from the ancestral function, which appears to be maintained by TRFL7.

### TRFL7/5/11 proteins localize to telomeres

We next assessed the intracellular localization of Arabidopsis TRFL7 and its homologs. Transient expression of N- and C-terminally YFP-fused TRFL7, TRFL5, and TRFL11 constructs in Arabidopsis mesophyll protoplasts resulted in a nuclear signal with prominent speckles (Figure 4A, Figure S3A). Co-transfection of the tagged TRFL7, TRFL5, and TRFL11 proteins showed a perfect overlap of the nuclear speckle signals (Figure 4B), indicating that these proteins colocalize to the same nuclear structures. Co-expression with nucleolar (fibrillarin) and Cajal body (coilin) markers showed a distinct pattern from the TRFL7 speckles, suggesting that these form different subnuclear compartments (Figure 4C). The prominent speckles were characteristic of TRFL7/5/11, as the other proteins of TRFL family 2 exhibited a diffuse nuclear signal or less prominent speckles (Figure S3B). This also applies to TRFL9, a representative of TRFL family 1, which displayed a diffuse nuclear signal (Figure S3B).

**Figure 4.**
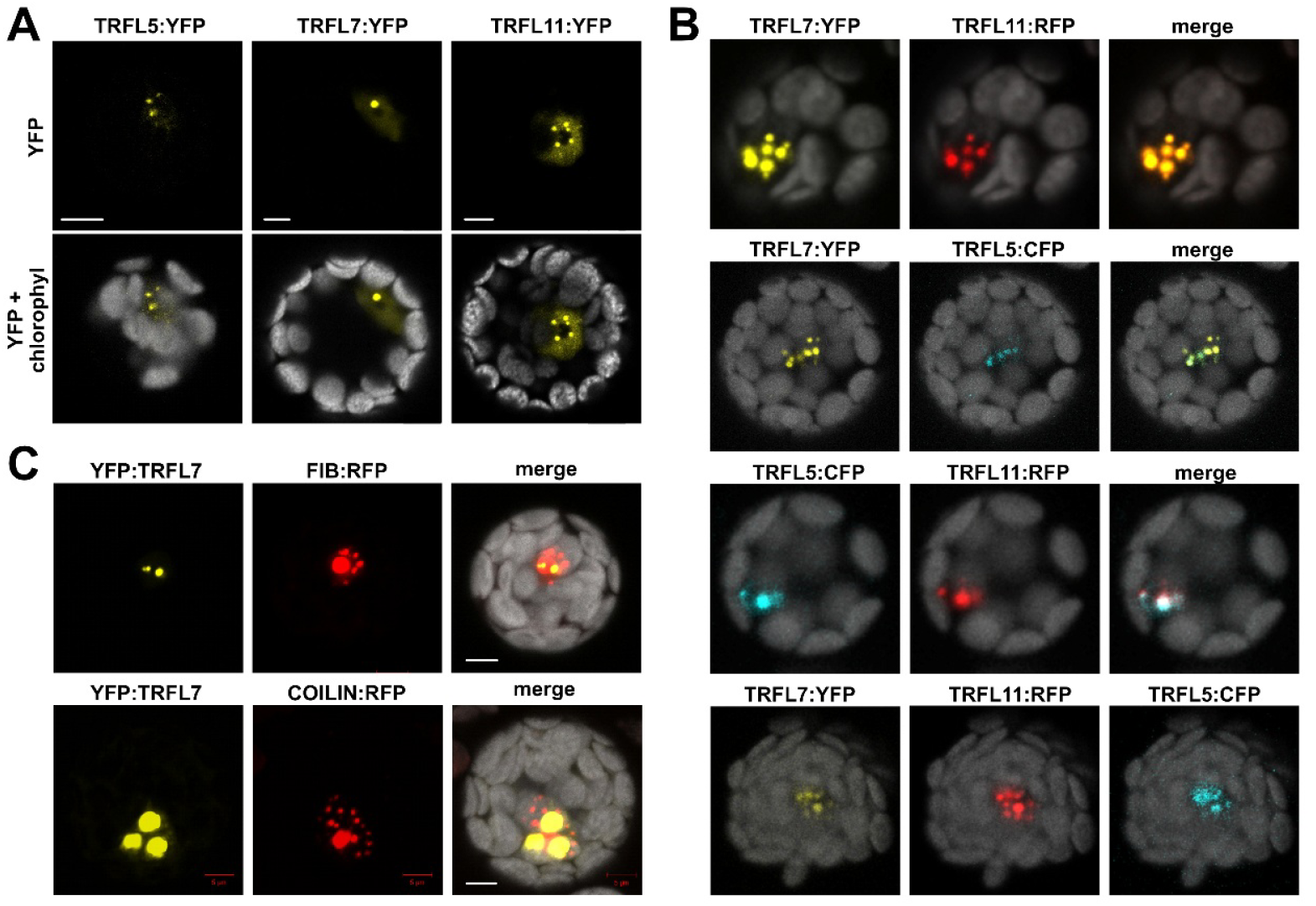
Localization of Arabidopsis TRFL homologues in mesophyll protoplasts. **(A)** Subcellular localization of TRFL5, TRFL7, and TRFL11 fused to YFP at the C terminus. (**B**) Co-localization of TRFL7 with nucleolar (fibrillarin, FIB) and Cajal body (coilin) markers. (**C**) Mutual co-localization of TRFL5, TRFL7, and TRFL11. Scale bar = 5 µm.

To further validate the localization data obtained from transiently transfected protoplasts, we generated Arabidopsis transgenic lines expressing C-terminally YFP-fused TRFLs under the control of the 35S promoter. The signal from TRFL7-YFP lines was too weak for reliable detection. Nevertheless, root cells in TRFL5-YFP and TRFL11-YFP lines produced a nuclear signal with clearly distinct foci (Figure 5A). In some cells, these foci formed a ring at the boundary of the nucleolus (Figure 5B,C). Three-dimensional analysis of mesophyll protoplasts from TRFL5-YFP plants transfected with a nucleolar marker confirmed that some of the TRFL5 speckles tend to associate with the nucleolar periphery. These foci are relatively uniform in size, although they occasionally form a larger granule (Figure 5D). Such a localization pattern is reminiscent of telomeres, which are roughly similar in size and typically organized around the nucleolus in Arabidopsis somatic and meiotic cells (59,60).

**Figure 5.**
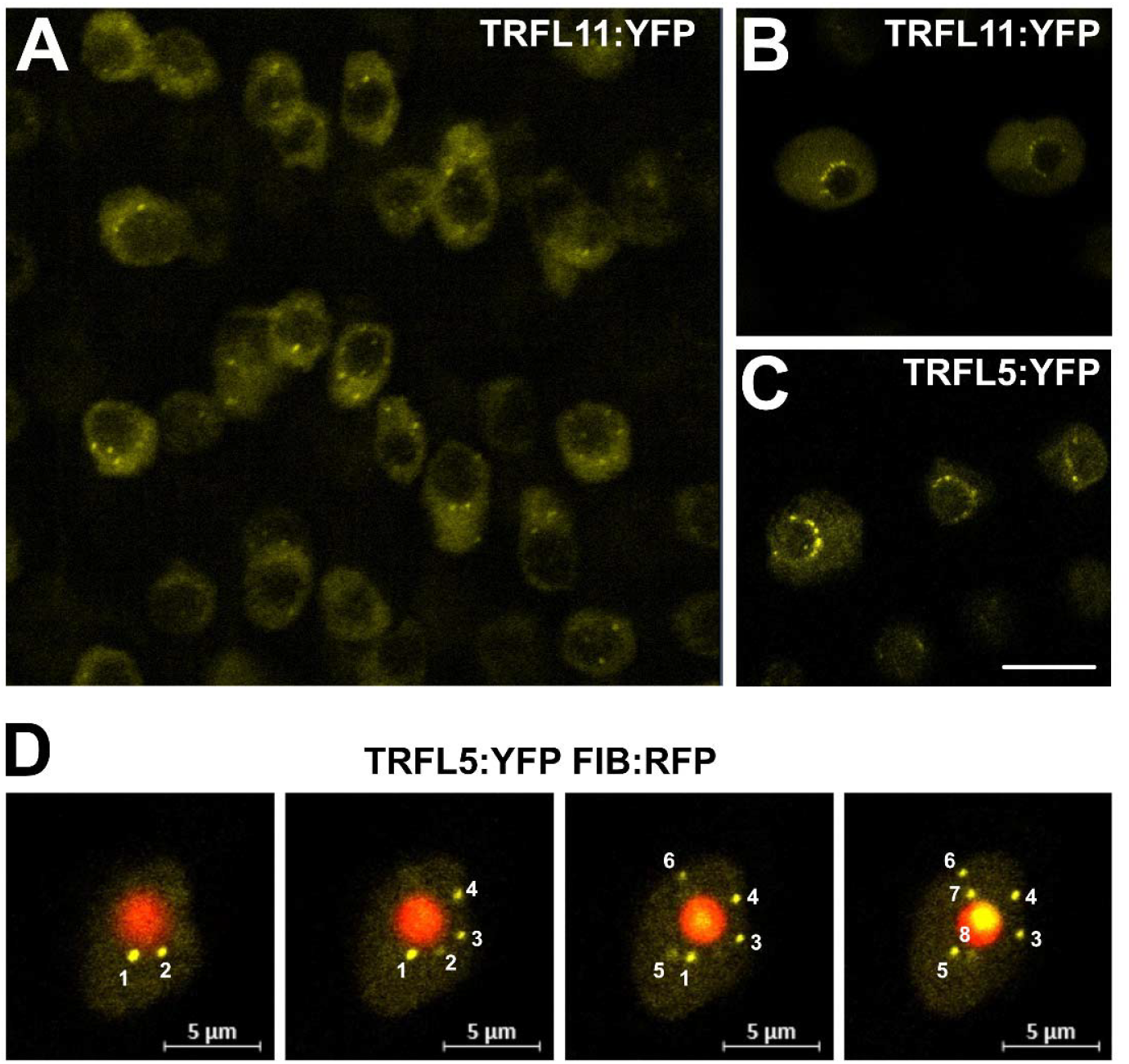
Localization of TRFL homologues in transgenic plants. **(A)** Micrograph showing a representative image of root cells expressing TRFL11:YFP. (**B, C**) Micrographs showing selected root cells expressing TRFL11:YFP and TRFL5:YFP, with speckles localized around the nucleolus. (**D**) Different Z-sections of the same mesophyll protoplast derived from a TRFL5:YFP plant that was transiently transfected with the FIB:RFP nucleolar marker. Individual speckles, as they appear in different Z-sections, are numbered. Scale bars = 5 µm.

We next performed chromatin immunoprecipitation of YFP-tagged proteins followed by qPCR to examine whether TRFL7/5/11 bind to telomeric chromatin. We found strong enrichment of telomeric DNA in the TRFL7-YFP, TRFL5-YFP, and TRFL11-YFP fractions immunoprecipitated with GFP antibody, as well as in histone H2A.10-GFP precipitates, compared to the wild-type (no YFP) control (Figure 6 and S4). In contrast, no enrichment of telomeric DNA was detected in pull-downs from plants expressing CDKD3- YFP, a protein localized in the nucleus (61). We found no enrichment of centromeric DNA in the immunoprecipitates from TRFL-YFP plants, although this signal was present in H2A.10-GFP samples. These results show that TRFL7/5/11 proteins specifically associate with telomeric chromatin, which, together with the microscopy data, provides strong evidence for their binding to telomeres.

**Figure 6.**
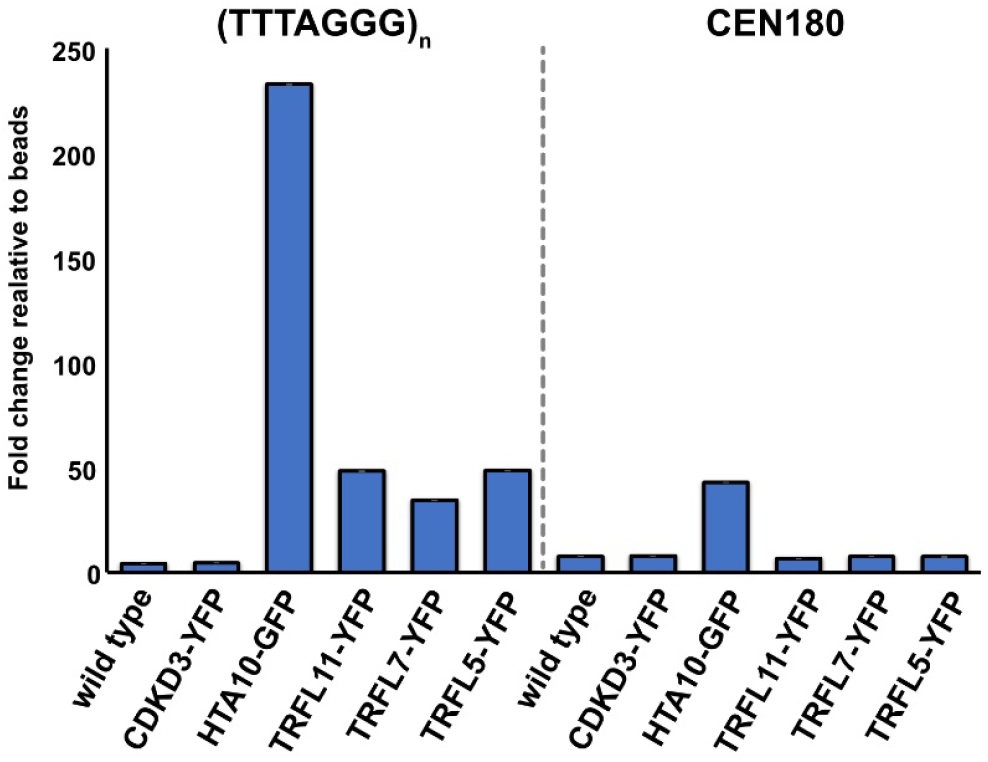
Detection of TRFL proteins in telomeric chromatin by chromatin immunoprecipitation. The bar chart indicates fold enrichment over beads lacking the anti-GFP antibody. Error bars represent the SD from three technical replicates. Data from an independent ChIP experiment are presented in Figure S4.

### TRFL7 forms homo- and heterodimers

Human TRF1 and TRF2 bind to telomeres as homodimers, and a similar capacity for homodimerization has also been observed among members of the Arabidopsis TRFL Family 1 (38). Therefore, we performed yeast two-hybrid assays to examine interactions among the TRFL7, TRFL5, and TRFL11 homologues. TRFL7 was capable of both self-interaction and interactions with TRFL5 and TRFL11 (Figure 7). In contrast, TRFL5 and TRFL11 exhibited only a weak propensity to heterodimerize with each other and lacked the ability to form homodimers (Figure 7). Structural studies of hTRF1 and hTRF2 indicate that two SANT/Myb domains are required for stable binding to telomeric DNA (62), suggesting that multimerization may also be important for efficient telomere association by Arabidopsis TRFL proteins. Thus, the ability of TRFL7 to homodimerize and interact with other members of this subfamily indicates an ancestral role in telomere-related functions.

**Figure 7.**
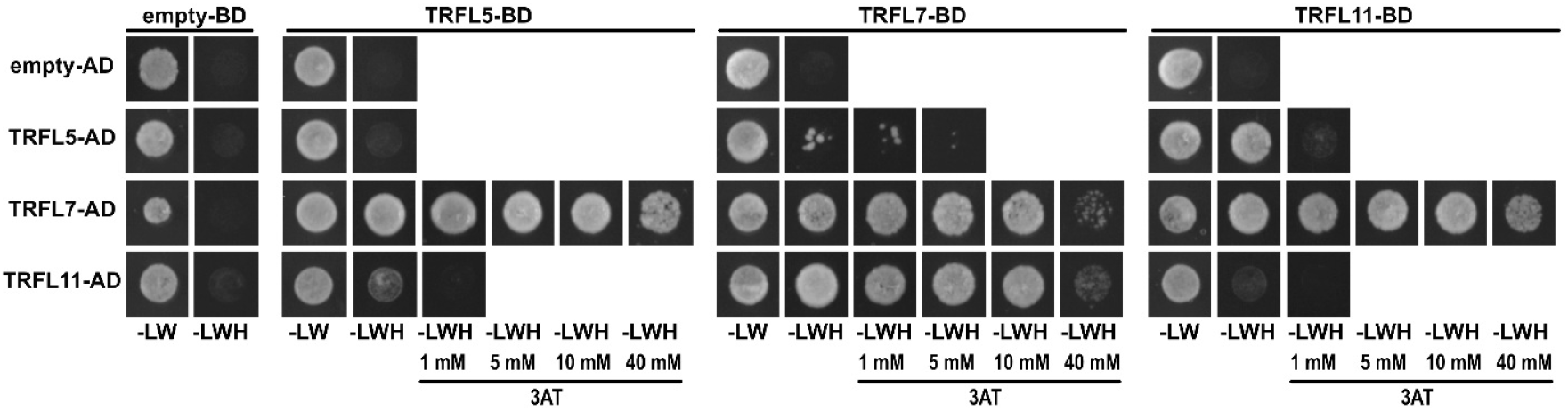
Interactions between the indicated TRFL homologues assessed by a yeast two-hybrid assay. Yeast cells expressing the indicated constructs were grown on either non-selective (-LW) or selective (-LWH) media supplemented with increasing concentrations of 3-amino-1,2,4-triazole (3-AT).

### Effect of TRFL7 and its homologues on telomere length

To investigate the impact of TFL7/5/11 proteins on telomeres, we analyzed Arabidopsis lines harboring T-DNA insertions disrupting the respective genes (Figure S5). Plants homozygous for these T-DNA insertions did not exhibit any obvious growth phenotypes and were fully fertile (Figure S6). Terminal restriction fragment analysis in individual mutant plants showed telomeres ranging from 2 to 4 kb (Figures 8 and S5), which is typical for wild-type plants.

**Figure 8.**
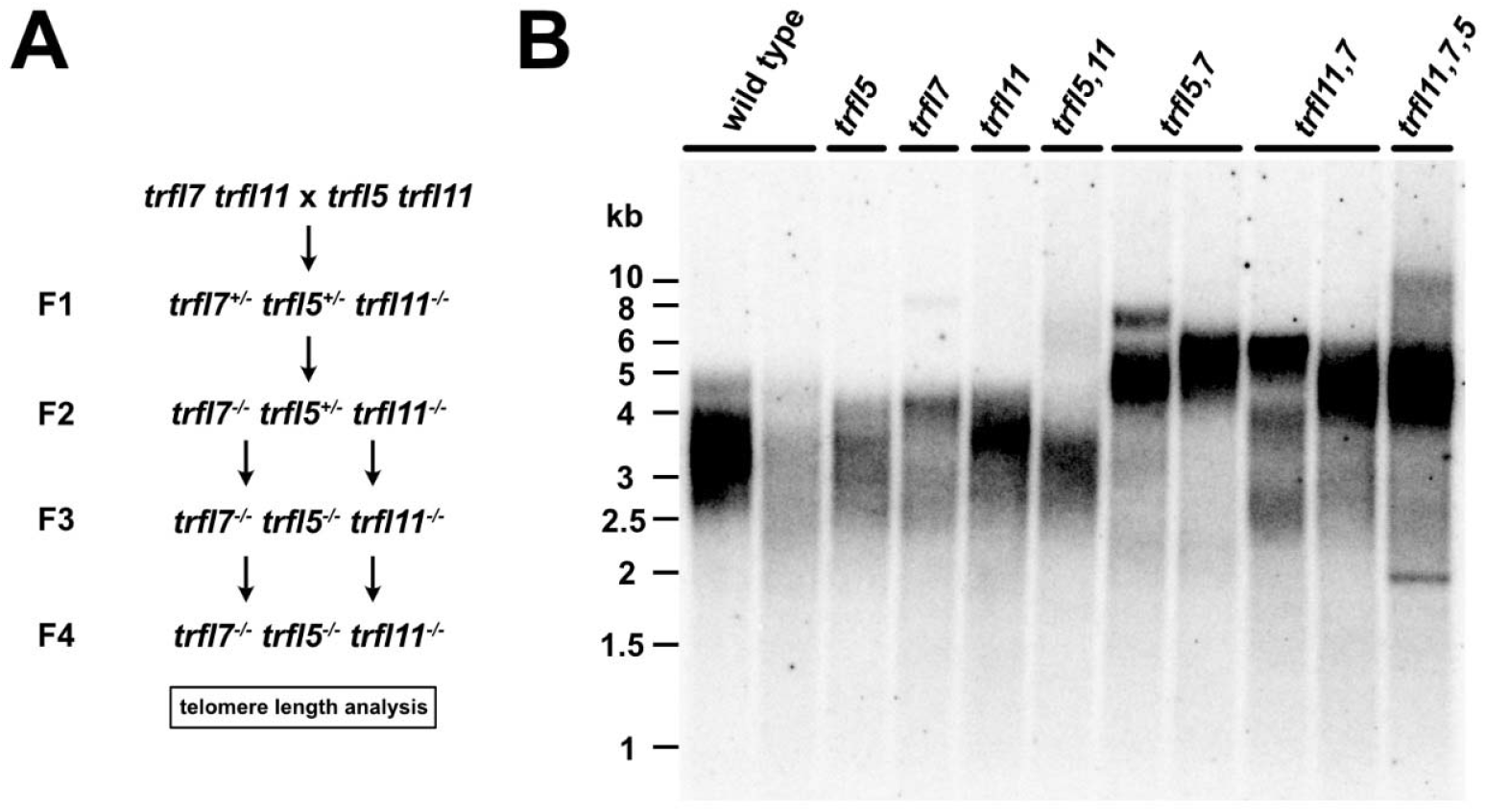
Terminal restriction fragment analysis in the indicated *trfl* mutants. **(A)** Crossing scheme used to generate *trfl7 trfl5 trfl11* triple mutants. (B) Terminal restriction fragment analysis of individual plants of the respective genotypes. Plants were analyzed after being propagated for two generations as homozygous mutants.

Since TRFL5/7/11 co-localize to telomeres and are of relatively recent evolutionary origin, they may carry partially redundant functions. To examine this possibility, we generated all combinations of double mutants, as well as a triple mutant line, by crossing *trfl5 trfl11*-1 with *trfl7 trfl11*-1 double mutants (Figure 8A). All combinations of mutations resulted in phenotypically normal, fertile plants (Figure S6). Nevertheless, telomere length analysis revealed that while *trfl5 trfl11*-1 mutants exhibited wild type-like telomeres ranging from 2 to 4 kb, telomeres were extended to up to 7 kb in *trfl7 trfl11*-1 and *trfl5 trfl7* double mutant combinations (Figures 8 and S7). Inactivation of all three homologs did not lead to further exacerbation of the telomeric phenotype.

Telomere length deregulation appeared to be progressive, as fourth-generation (G4) selfed double mutants exhibited longer telomeres than those observed in the second generation (G2) (Figure S7). This pattern suggests a gradual shift in telomere length homeostasis rather than an abrupt extension. Together, these findings indicate partial functional redundancy among the TRFL homologs and support a central role for TRFL7, as only mutant combinations that include *trfl7* display telomere length deregulation.

## Discussion

While numerous proteins have been linked to telomere function and maintenance in Arabidopsis, the factors that specifically bind to telomeric duplex DNA and are expected to confer the unique functional properties of telomeric chromatin have remained elusive. Here, we used telomeric DNA affinity purification followed by mass spectrometry with the aim to identify the full complement of Arabidopsis telomere-binding proteins. We found TRFL7 and TRB3 among the most highly enriched telomeric DNA– binding proteins. Arabidopsis TRBs and their rice homologs have previously been identified using similar proteomic approaches, further validating the association of these proteins with plant telomeres (35,63). TRFL7 was the only member of the TRFL protein family enriched in the telomeric DNA–bound fraction. Importantly, the TRFL7 homolog in rice, known as ANTHER INDEHISCENCE1, was also identified as the sole TRFL representative in telomeric DNA pull-downs (63), suggesting a more prominent telomere-related role than other members of this gene family.

Several additional evidence support the notion that TRFL7 is an important functional constituent of telomeric chromatin. First, subcellular localization analyses show that TRFL7 and its homologs, TRFL5 and TRFL11, form prominent nuclear speckles that tend to associate with the nucleolar periphery and, in some cells, even align around the nucleolus in a necklace-like pattern. Such an arrangement is typical of telomere organization in Arabidopsis interphase nuclei (59,60), and hence, these speckles may represent telomeres. Second, we found that TRFL7 and its homologs specifically bind to telomeric chromatin, as demonstrated by chromatin immunoprecipitation. Third, we show that Arabidopsis mutants deficient in TRFL7 in combination with either TRFL5 or TRFL11 exhibit telomere elongation. Finally, phylogenetic analysis suggests that plant TRFL7 and the metazoan TRF2 proteins share a common evolutionary origin and potentially related functionality, as they display similar protein topology and are under selective constraint to retain the iDDR domain.

The iDDR has been identified in human TRF2 as a sequence motif that inhibits activation of ATM at telomeres and prevents their resection by the Mre11–Nbs1–Rad50 (MRN) complex (58,64). The iDDR exerts its function by binding to Rad50 and modulating the activity of the MRN complex, presumably by inhibiting its association with CtIP and DNA (64,65). The iDDR motif was initially thought to be specific to metazoan TRF2-like proteins. However, functionally analogous, though sequence-unrelated, motifs termed MIN (for MRN inhibitor) have also been identified in Rif2 and Taz1, key constituents of telomeric chromatin in budding and fission yeasts (66,67). Thus, although the evolutionary origins of yeast and animal telomere-binding proteins are distinct, they appear to have converged on a similar functional strategy involving direct interaction with Rad50 to restrain MRN activity.

The discovery of an iDDR-like motif in plant TRFL7 homologs provides a new perspective on the origin of telomere-binding proteins and the mechanisms of chromosome-end protection. The presence of TRFL7- like proteins in both streptophytic and chlorophytic green algae, which diverged near the base of the plant kingdom approximately one billion years ago, suggests that these proteins reflect an ancient structural form of telobox proteins. The similarity in protein topology between plant TRFL7 and animal TRF2 further supports their origin in the last eukaryotic common ancestor, and the selective constraint on the preservation of the iDDR motif suggests that modulation of the MRN complex represents a central and ancient principle of chromosome-end protection.

However, it remains to be experimentally established whether plant iDDR indeed interacts with the MRN complex and modulates its activity at telomeres. While TRFL7 and TRF2 are similar in their topology, ability to form dimers, and capacity to interact with telomeric DNA through the SANT/Myb domain, the impact of TRFL7 on chromosome-end protection appears to be relatively mild compared to human TRF2, the inactivation of which induces massive chromosome end-to-end fusions. In Arabidopsis, telomere fusions triggered either by telomere depletion due to the absence of telomerase or by inactivation of the CST complex result in ongoing genome instability that impairs plant growth and fertility (19,20,68,69). None of these phenotypes have been observed in Arabidopsis mutants deficient in all three TRFL7 paralogues.

So far, the only discernible telomere-related phenotype in these mutants is telomere elongation. This elongation is likely mediated by telomerase, suggesting an inhibitory role for TRFL7 in regulating telomere synthesis. Thus, it is possible that any chromosome-end capping defects are compensated by telomerase-mediated telomere extension. A similar situation occurs in Ku-deficient plants, which exhibit exonuclease-mediated telomere degradation and increased recombination, yet maintain functional telomeres through telomerase-dependent elongation (16,70). Accelerated telomere deprotection is unmasked only in plants doubly deficient in Ku and telomerase (71).

The relatively weak telomere-related phenotypes observed in the most extensively characterized plant telobox protein candidates suggest that either an essential telomere-binding protein required for chromosome-end protection remains unidentified, or this function is shared among several proteins that collectively ensure end protection while being individually largely dispensable. In humans, shelterin appears to exist in distinct subcomplexes that provide flexibility to accommodate different telomere maintenance functions (72,73).

Such specialization may also exist in Arabidopsis, where at least two classes of telomere-binding proteins, TRBs and TRFL7, localize to telomeres. TRFL7, which appears to be more closely related to a putative ancient shelterin-like complex, may be in Arabidopsis more specialized in telomerase regulation. Notably, another conserved shelterin component, POT1B, is required in Arabidopsis to stimulate telomerase activity at telomeres but not for chromosome-end protection (22,74). The complexity of protein organization and functional interactions at plant telomeric chromatin is further increased by the presence of multiple gene family members, which likely adds robustness and redundancy to the system. Identification of TRFL7 as a putative TRF2 homologue provides new impetus for future studies into the organization and regulation of plant telomeres, as well as the evolution of mechanisms of telomere maintenance and protection.

## Acknowledgement

We acknowledge the support from CEITEC MU Core facilities Plant Sciences, CELLIM Facility of Czech-Bioimaging supported by MEYS CZ (No. LM2018129), and Proteomics Core Facility of CIISB, Instruct-CZ Centre, supported by MEYS CR (LM2023042, CZ.02.01.01/00/23_015/0008175, e-INFRA CZ (ID:90254). This work was funded by the Czech Science Foundation (grant 21-25163J to KR).

## Notes

### Competing Interest Statement

The authors have declared no competing interest.

