## Supplementary material for "Identification and characterization of a TRF2-like telomere-binding protein in Arabidopsis": Figures S1-S7

**Supplementary information**

Supplementary Figures S1- S6

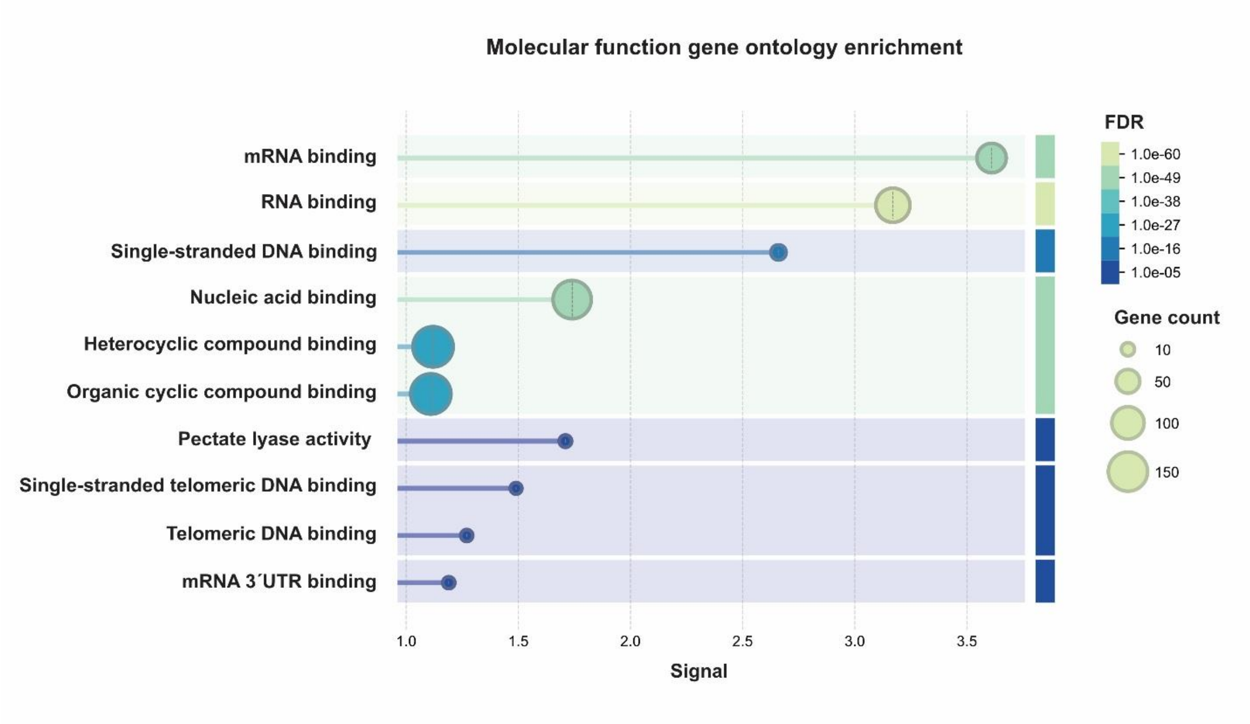


**Figure S1.** Molecular function gene ontology enrichment analysis of proteins specifically co-purifed with the 50 bp telomeric probe.

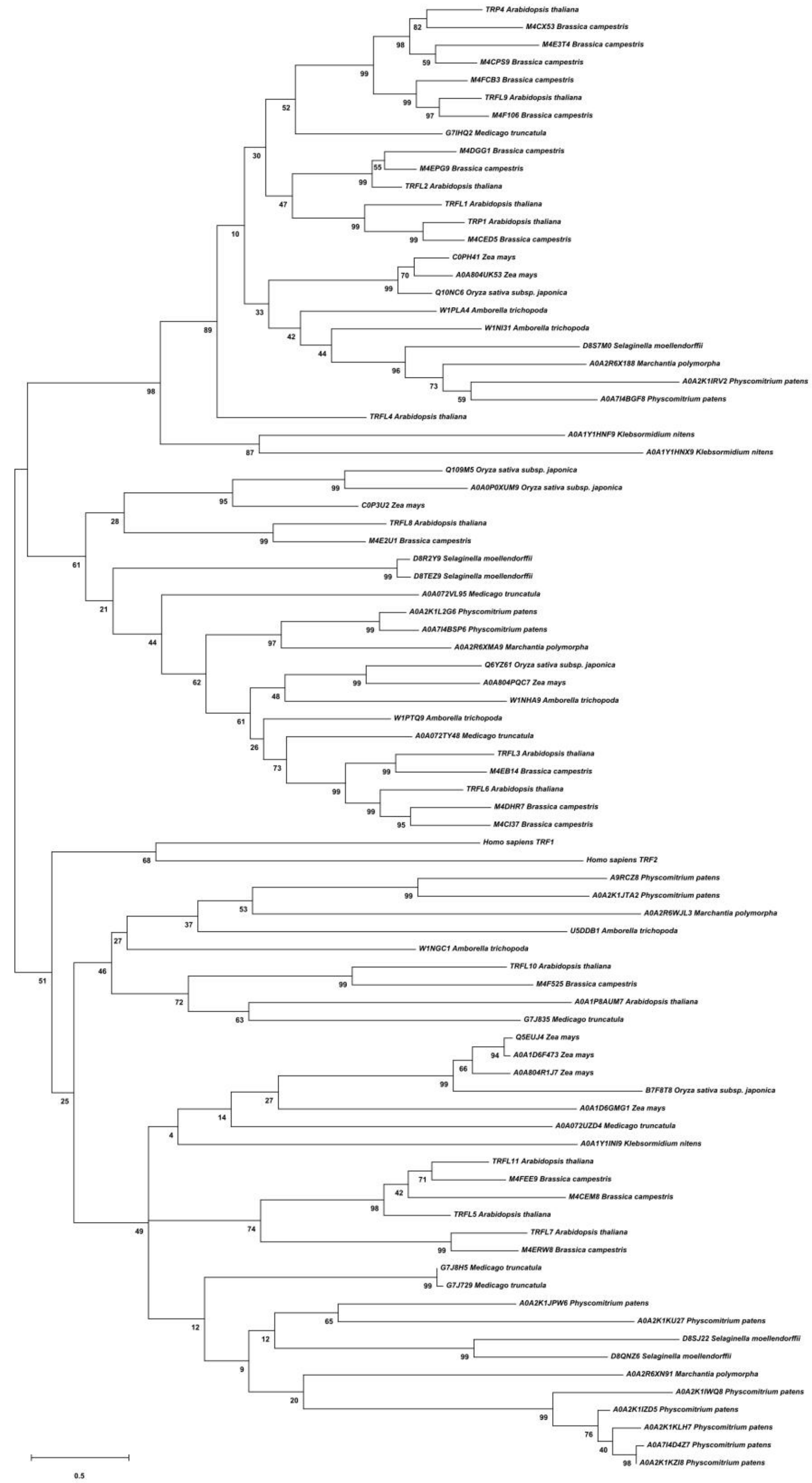

**Figure S2.** Phylogenetic tree of plant TRFL homologues. The bootstrap values and UniProt protein identification numbers are indicated. Scale bar indicates evolutionary distance.

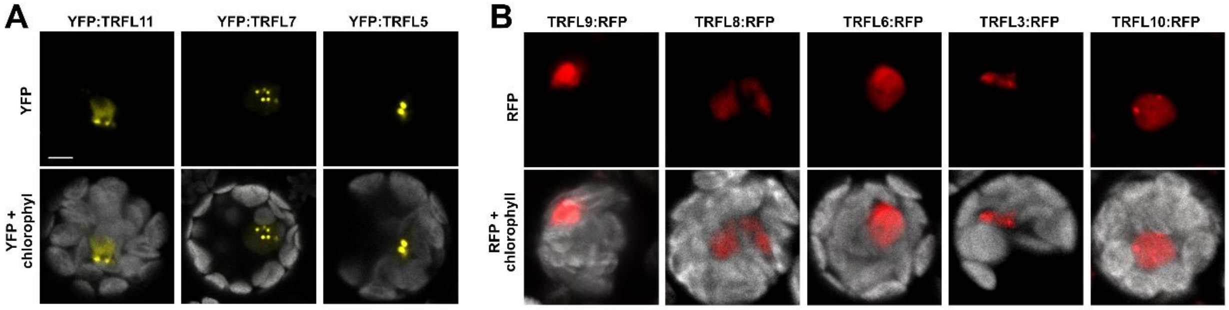


**Figure S3.** Localization of Arabidopsis TRFL homologues in mesophyll protoplasts. (**A**) Subcellular localization of TRFL5, TRFL7, and TRFL11 fused to YFP at the N terminus. (**B**) Localization of TRFL representatives from other TRFL7 subgroups as indicated in Figure S3. Scale bar = 5 µm.

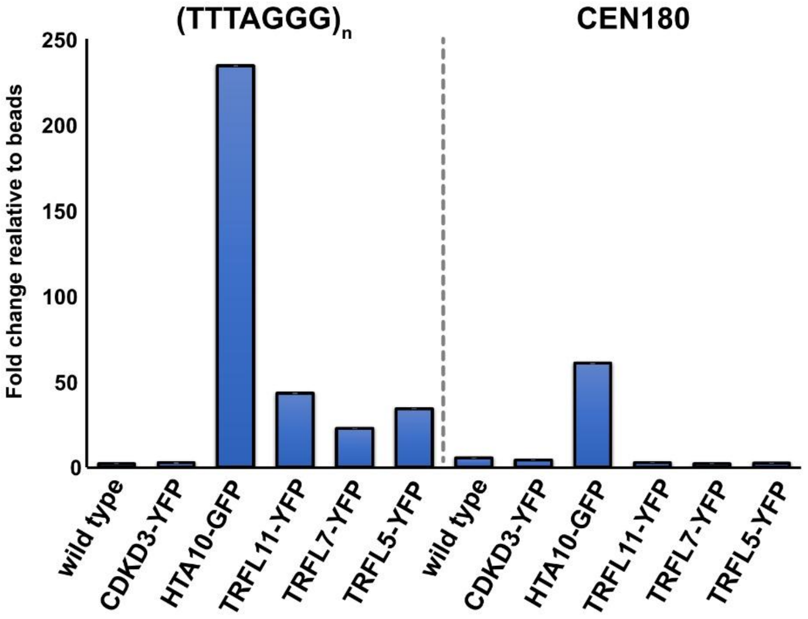


**Figure S4.** Detection of TRFL proteins in telomeric chromatin by chromatin immunoprecipitation. Independent replica of the experiment shown in Figure 6. Error bars represent the SD from three technical replicates.

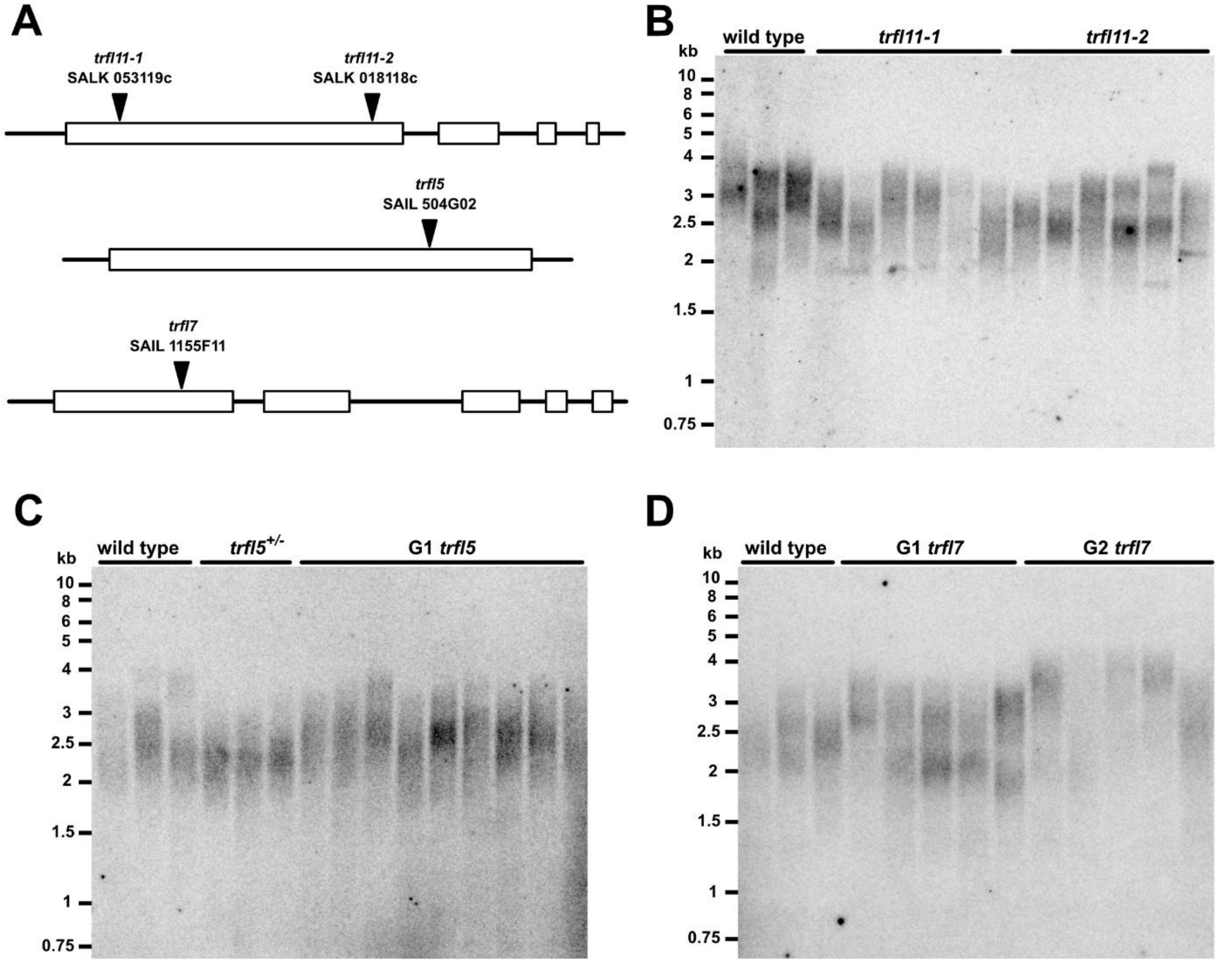


**Figure S5.** Terminal restriction fragment analysis in *trfl* single mutants. (**A**) Scheme of the T-DNA insertion lines used in this study. Diagrams of the *TRFL11*, *TRFL7*, and *TRFL5* genes are shown, with exons depicted as rectangles. T-DNA insertions are indicated. (**B–D**) Terminal restriction fragment analysis of the indicated mutants. First-generation mutants were analyzed for *trfl5*, whereas both first- and second-generation mutants were analyzed for *trfl7*. The *trfl11-1* and *trfl11-2* lines were obtained as confirmed homozygous mutants at an unknown generation, as the exact generation after out-segregation from the heterozygous parental line could not be determined.

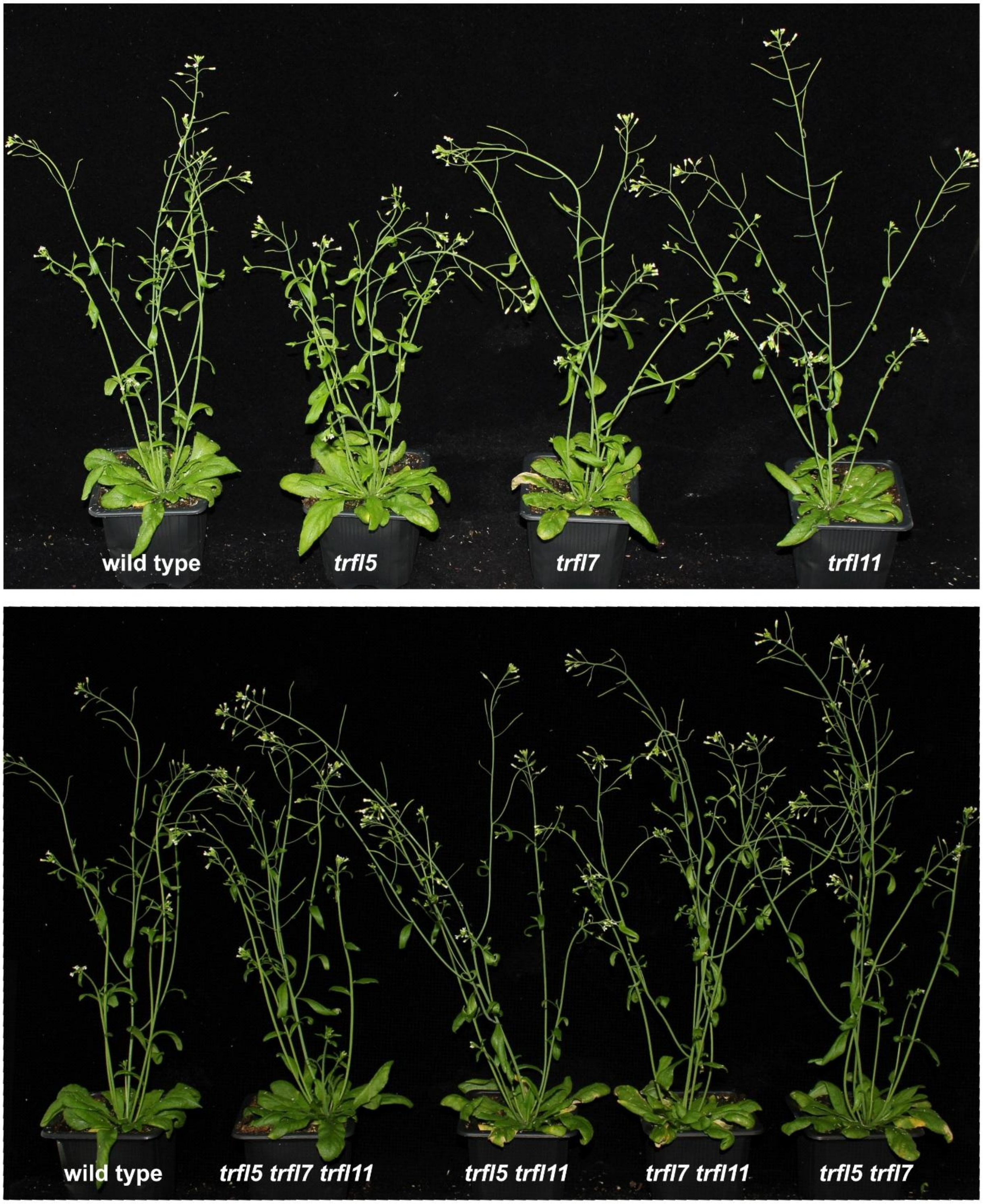


**Figure S6.** Pictures of *trfl* mutant plants.

In the

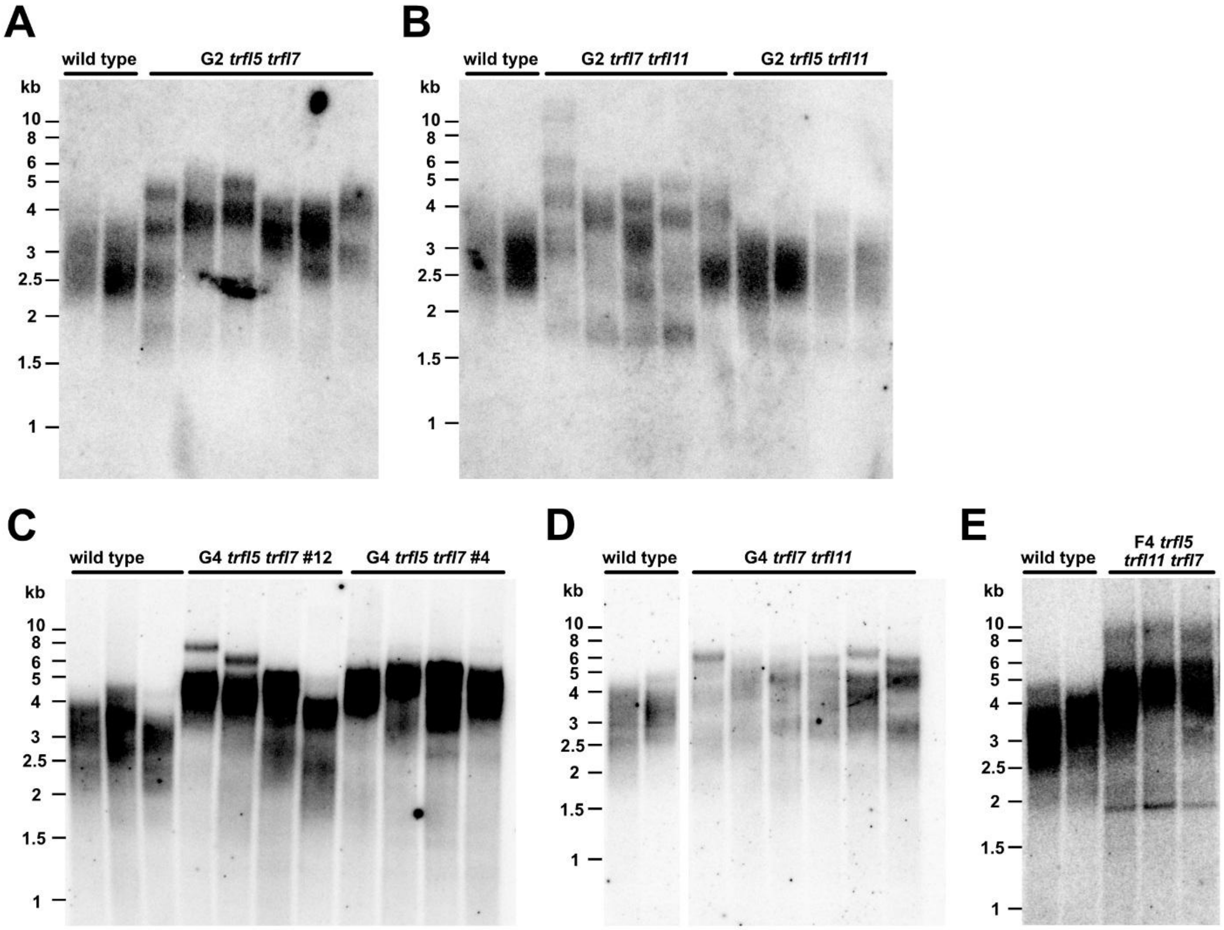


**Figure S7.** Terminal restriction fragment analysis in the indicated double-mutant combinations in the second (**A, B**) and fourth (**C, D**) generations. (**E**) Telomere length in the triple mutants in F4 (second-generation triple mutants; see Figure 8A).
